# Continual inactivation of genes involved in stem cell functional identity stabilizes progenitor commitment

**DOI:** 10.1101/2020.02.03.931972

**Authors:** Noemi Rives-Quinto, Hideyuki Komori, Derek H. Janssens, Shu Kondo, Qi Dai, Adrian W. Moore, Cheng-Yu Lee

## Abstract

Expansion of the pool of stem cells that indirectly generate differentiated cells through intermediate progenitors drives vertebrate brain evolution. Due to a lack of lineage information, mechanistic investigation of the competency of stem cells to generate intermediate progenitors remains impossible. Fly larval brain neuroblasts provide excellent *in vivo* models for investigating the regulation of stem cell functionality during neurogenesis. Type II neuroblasts undergo indirect neurogenesis by dividing asymmetrically to generate a neuroblast and a progeny that commits to an intermediate progenitor (INP) identity. We identified Tailless (Tll) as the master regulator that maintains type II neuroblast functional identity, including the competency to generate INPs. Successive inactivation during INP commitment inhibits *tll* activation by Notch, preventing INPs from reacquiring neuroblast functionality. We propose that the continual inactivation of neural stem cell functional identity genes by histone deacetylation allows intermediate progenitors to stably commit to generating diverse differentiated cells during indirect neurogenesis.

## Introduction

The outer subventricular zone (OSVZ) is unique to gyrencephalic mammals and directly contributes to the immense disparity observed in the appearance and functionality of their brains compared with those of lissencephalic mammals (Cárdenas and Borrell, 2019; Delaunay et al., 2017; Di Lullo and Kriegstein, 2017). A key functional feature of OSVZ neural stem cells is their dependence on intermediate progenitors to indirectly generate neurons (Cárdenas and Borrell, 2019; Delaunay et al., 2017; Di Lullo and Kriegstein, 2017). Intermediate progenitors function as transit-amplifying cells to increase cell number and diversity, allowing for the formation of deep folding, which is characteristic of gyrencephalic brains. Despite recent advances in our understanding of the generation of OSVZ neural stem cells (Fujita et al., 2019; Namba et al., 2019), key open questions regarding the mechanisms regulating their unique functionality, such as the competency to generate intermediate progenitors remain.

Two distinct neuroblast lineages function together to generate the number of neurons requisite for the development of adult fly brains (Farnsworth and Doe, 2017; Homem et al., 2015; Janssens and Lee, 2014). Similar to ventricular zone neural stem cells, type I neuroblasts undergo direct neurogenesis. Type I neuroblasts repeatedly undergo asymmetric division to generate one daughter cell that remains a neuroblast and one sibling cell [ganglion mother cell (GMC)] that divides once to generate two neurons. By contrast, type II neuroblasts undergo indirect neurogenesis, similar to OSVZ neural stem cells. Type II neuroblasts continually undergo asymmetric division to self-renew and to generate a sibling cell that commits to an INP identity (Bello et al., 2008; Boone and Doe, 2008; Bowman et al., 2008). The Sp8 family transcription factor Buttonhead (Btd) and the ETS-1 transcription factor PointedP1 (PntP1) are specifically expressed in type II neuroblasts (Komori et al., 2014a; Xie et al., 2014; Zhu et al., 2011). Btd and PntP1 expression levels decrease as a newly generated immature INP transitions into a non-Asense-expressing (Ase^−^) immature INP. Three to four hours after this transition, an Ase^−^ immature INP upregulates Ase expression as it progresses through INP commitment. Once INP commitment is complete, an Ase^+^ immature INP transitions into an INP that undergoes 6-8 rounds of asymmetric division. These molecularly defined intermediate stages during INP commitment provide critical landmarks to identify genes that control type II neuroblast functional identity.

Notch signaling maintains type II neuroblasts in an undifferentiated state, partially by poising transcription of the master regulator of differentiation *earmuff* (*erm*) (Janssens et al., 2017). During asymmetric neuroblast division, the TRIM-NHL protein Brain tumor (Brat) segregates into the newly generated immature INP and targets transcripts encoded by Notch downstream effector genes for RNA decay (Bello et al., 2006; Betschinger et al., 2006; Komori et al., 2018; Lee et al., 2006; Xiao et al., 2012). *brat-*null brains accumulate thousands of supernumerary type II neuroblasts that originate from the reversion of newly generated immature INPs due to defects in the downregulation of Notch signaling (Komori et al., 2014b). In parallel to Brat, the nuclear protein Insensible (Insb) inhibits the activity of Notch downstream effector proteins during asymmetric neuroblast division, and Insb overexpression efficiently triggers premature differentiation in type II neuroblasts (Komori et al., 2018). This multi-layered gene control allows for the onset of Erm expression, coinciding with the termination of PntP1 expression. Erm expression is maintained in immature INPs but rapidly diminishes in INPs (Janssens et al., 2014). Erm belongs to a family of transcription factors that are highly expressed in neural progenitors and can bind histone deacetylase 3 (Hdac3) (Hirata et al., 2006; Koe et al., 2014; Levkowitz et al., 2003; Weng et al., 2010). Additionally, Erm prevents INP reversion by repressing gene transcription. In *erm*-null brains, INPs spontaneously revert to type II neuroblasts; however, this phenotype can be suppressed by knocking down *Notch* function (Weng et al., 2010). Thus, Erm likely inactivates type II neuroblast functionality genes by promoting histone deacetylation.

By comparing mRNAs enriched in type II neuroblasts or immature INPs, we identified *tailless* (*tll*) as the master regulator of type II neuroblast functional identity. Tll is expressed in type II neuroblasts but not in INPs; moreover, Tll is necessary and sufficient for type II neuroblast functionality. Tll overexpression is sufficient to transform type I neuroblasts into type II neuroblasts, as indicated by changes in gene expression and an acquired competence for generating INPs. We identified *hamlet* (*ham)* as a new negative regulator of type II neuroblast maintenance. *ham* is expressed after *erm*, and Erm and Ham function through Hdac3 to continually inactivate *tll*. Sequential inactivation during INP commitment suppresses *tll* activation by Notch signaling in INPs. We propose that continual inactivation of the master regulator of stem cell functional identity by histone deacetylation allows intermediate progenitors to stably commit to the generation of differentiated cell types without reacquiring stem cell functionality.

## Results

### A novel transient over-expression strategy to identify regulators of type II neuroblast functional identity

Genes that regulate neuroblast functional identity should be expressed in type II neuroblasts and become rapidly downregulated in Ase^−^ and Ase^+^ immature INPs in wild-type brains (Figure 1A). To enrich for transcripts that fulfill these criteria, we tested if supernumerary type II neuroblasts transiently overexpressing a *UAS-insb* transgene synchronously transition into INPs in *brat-*null brains as in wild-type brains. The combination of *Wor-Gal4, AseGal80*, which is only active in type II neuroblasts, and *Tub-Gal80*^*ts*^, which loses its inhibitory effect on Gal4 activity under non-permissive temperatures, allows for spatial and temporal control of Insb overexpression in all type II neuroblasts. We allowed larvae to grow at 25°C, and transiently overexpressed Insb for 6, 12, 18, or 24 hours by shifting the larvae to a non-permissive temperature of 33°C. Larvae that remained at 25°C for the duration of this experiment served as the control (time 0). We assessed the effect of this transient over-expression strategy on cell identity by quantifying the total type II neuroblasts and INPs per brain lobe based on Deadpan (Dpn) and Ase expression (Figure 1A). Overexpressing Insb for 6 hours did not significantly affect the supernumerary neuroblast phenotype in *brat-*null brains (Figure S1). By contrast, 12 or more hours of Insb overexpression led to a time-dependent decrease in supernumerary type II neuroblasts and a corresponding increase in INPs. These results demonstrate that the transient Insb overexpression can induce supernumerary type II neuroblasts to synchronously transition into INPs in *brat-*null brains.

**Figure 1.**
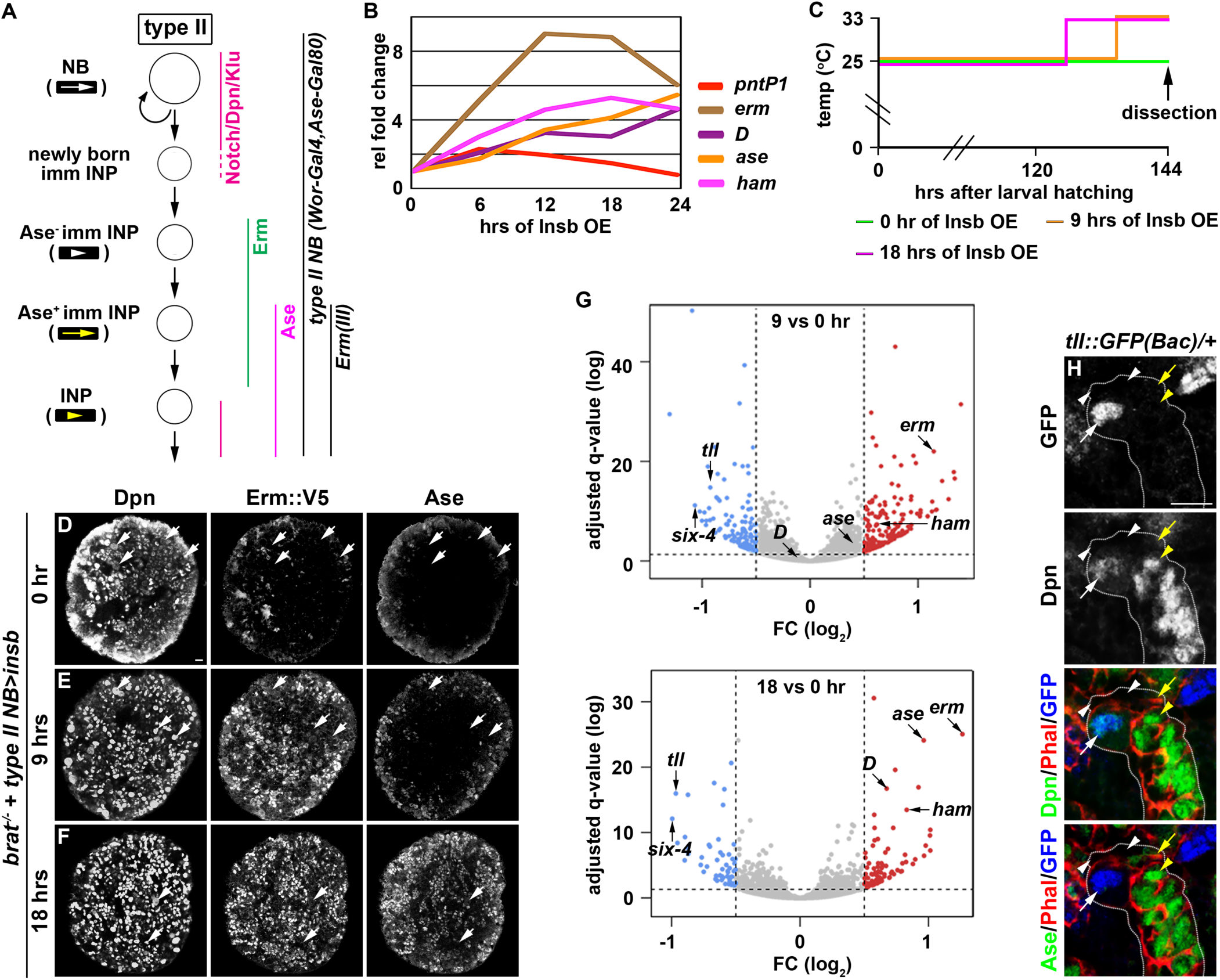
Identification of candidate regulators of type II neuroblast functional identity. (A) A diagram of the type II neuroblast lineage showing the expression patterns of genes and *Gal4* drivers. The color scheme of arrows and arrowheads used to identify various cell types in the type II neuroblast lineage in all figures is shown. (B) A time course analysis of gene transcription patterns in *brat*-null brains following transient Insb overexpression. Transient Insb overexpression triggers supernumerary type II neuroblasts in *brat*-null brains to activate gene transcription patterns that mimic immature INPs undergoing INP commitment in wild-type brains. (C) A strategy to synchronize the identities of supernumerary type II neuroblasts in *brat*-null brains to undergo INP commitment induced by transient Insb overexpression. Larvae were collected and aged at 25°C. Transient Insb overexpression was induced by shifting one third of larvae to 33°C at 126 or 135 hours after hatching. The last one third remained at 25°C and served as the source enriched for type II neuroblast-specific transcripts (time 0). (D-F) Confocal images of *brat*-null brains with transient Insb overexpression for 0, 9, or 18 hours. Transient Insb overexpression triggers supernumerary type II neuroblasts in *brat*-null brains to sequentially activate Erm and then Ase expression, recapitulating changes in marker expression during INP commitment in wild-type brains. Larvae used in this experiment carry the *erm::V5* allele generated by CRISPR-CAS9, allowing for the detection of endogenous Erm expression. (G) Volcano plots showing fold-change of gene expression in *brat*-null brains with transient Insb overexpression for 9 or 18 hours. (H) Tll expression pattern in the type II neuroblast lineage. Endogenous Tll is detected in type II neuroblasts but not in immature INPs and INPs in larval brains carrying one copy of the *tll::GFP(Bac)* transgene. Scale bar, 10 μm. See also Figure S1.

We next assessed if supernumerary type II neuroblasts transiently overexpressing Insb transition through identical intermediate stages to become INPs in *brat-*null brains as they do in wild-type brains (Figure 1A). We found that *pntP1* mRNA levels increased within the first 6 hours of Insb overexpression, and then continually declined (Figure 1B). By contrast, *erm* transcription rapidly increased in the first 12 hours of Insb overexpression, plateauing between 12 and 18 hours. *ase* transcription showed little change in the first 6 hours of Insb overexpression, and then steadily increased between 12 and 24 hours. The combined temporal patterns of *pntP1, erm*, and *ase* transcription strongly suggest that type II neuroblasts transiently overexpressing Insb for 6 hours in *brat-*null brains are at an equivalent stage as a newly generated immature INP transitioning to an Ase^−^ immature INP in wild-type brains. *brat-*null type II neuroblasts are at a stage equivalent to (1) Ase^−^ immature INPs following 6-12 hours of Insb overexpression and (2) Ase^+^ immature INPs following 12-24 hours of Insb overexpression. Indeed, following 9 hours of Insb overexpression most type II neuroblasts in *brat-*null brains expressed cell identity markers indicative of Ase^−^ immature INPs (Erm::V5^+^Ase^−^), whereas after 18 hours of Insb overexpression they expressed markers indicative of Ase^+^ immature INPs (Erm::V5^+^Ase^+^) (Figures 1C-F). These data indicate that supernumerary type II neuroblasts transiently overexpressing Insb indeed transition through identical intermediate stages during INP commitment in *brat-*null brains as in wild-type brains.

We predicted that a candidate regulator of type II neuroblast functional identity should become downregulated in *brat-*null brains following 9 and 18 hours of Insb overexpression. By sequencing mRNA in triplicate following the transient over-expression strategy (Figure 1C), we identified 76 genes that were reproducibly downregulated by 1.5-fold or more in *brat-*null brains overexpressing Insb for 9 hours (Figure 1G). Of these genes, *tll* was the most downregulated; similarly *tll* was the most downregulated gene in *brat-*null brains overexpressing Insb for 18 hours. We validated the *tll* expression pattern in the type II neuroblast lineage using a bacterial artificial chromosome (BAC) transgene [*tll::GFP(BAC)*] where green fluorescent protein (GFP) is fused in frame with the *tll* reading frame. Consistent with a previous study (Bayraktar and Doe, 2013), we detected Tll::GFP in type II neuroblasts but not in immature or mature INPs (Figure 1H). Thus, *tll* is uniquely expressed in type II neuroblasts, and is an excellent candidate for regulating type II neuroblast functionality.

### *tll* is the master regulator of type II neuroblast functional identity

We defined the type II neuroblast functional identity as the maintenance of an undifferentiated state and the competency to generate INPs. We first tested whether *tll* is required for maintaining type II neuroblasts in an undifferentiated state by overexpressing a *UAS-tll*^*RNAi*^ transgene to knock down *tll* function in larval brains. Whereas control brains always contained 8 type II neuroblasts per lobe, brains with *tll* function knocked down contained 4.3 ± 1.2 type II neuroblasts per lobe (Figure 2B; n = 10 brains per genotype). A closer examination revealed that the remaining *tll* mutant neuroblasts showed reduced cell diameters and ectopically expressed Ase, two characteristics typically associated with INPs in the type II neuroblast lineage (Figures 2A, 2C, 2D). This result indicated that *tll* is required for maintaining type II neuroblasts in an undifferentiated state. We next tested whether *tll* is sufficient to re-establish a type II neuroblast-like undifferentiated state in INPs by misexpressing a *UAS-tll* transgene under the control of the *Erm-Gal4(III)* driver (Figure 1A). Brains with *tll* misexpressed in INPs contained 150 ± 50 type II neuroblasts per lobe, whereas control brains always contained 8 type II neuroblasts per lobe (Figures 2E and 2F; Figure S2A; n = 10 brains per genotype). Thus, Tll misexpression is sufficient to revert INPs to type II neuroblasts. These data together led us to conclude that *tll* is necessary and sufficient for maintaining type II neuroblasts in an undifferentiated state.

**Figure 2.**
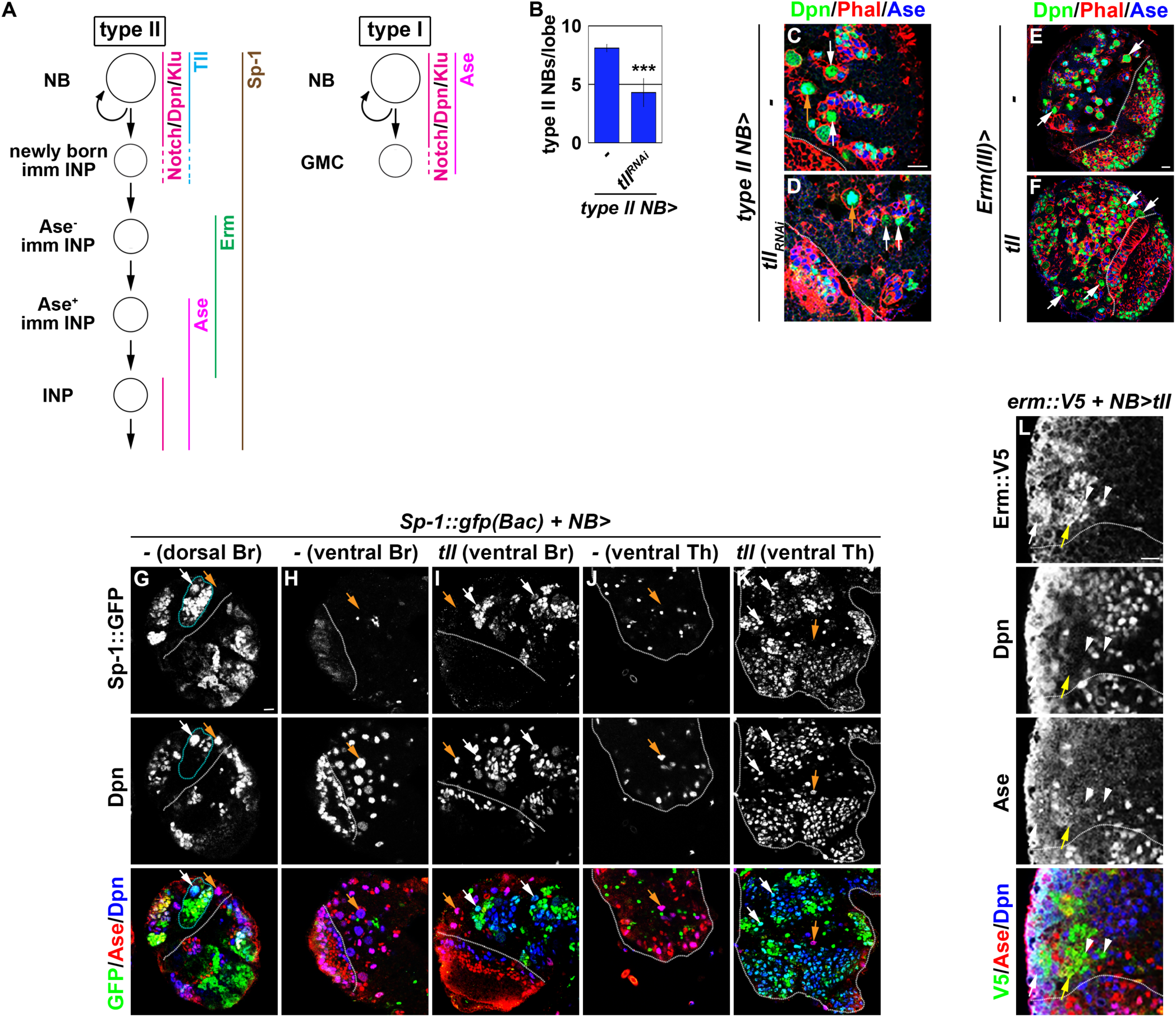
*tll* is necessary and sufficient for type II neuroblast functional identity. (A) A diagram of the type I and II neuroblast lineages showing the expression patterns of genes. (B) Quantification of total type II neuroblasts per lobe in larval brains with knockdown of *tll* function. Knocking down *tll* function reduced the number of type II neuroblasts. (C-D) Knocking down *tll* function led to premature differentiation in type II neuroblasts as indicated by reduced cell diameter and ectopic Ase expression (white arrows). Type I neuroblasts (orange arrows) served as the control. (E-F) Confocal images of larval brains with *tll* ectopically expressed in Ase^+^ immature INPs. *tll* mis-expression in Ase^+^ immature INPs led to supernumerary type II neuroblast formation. (G-H) Sp-1::GFP marks type II neuroblast lineages. Type I neuroblast lineages (orange arrows) were found on the dorsal and ventral surface of the brain (Br), but type II neuroblast lineages (white arrows) were only found on the dorsal surface. (I) Tll overexpression molecularly transforms type I neuroblasts into type II neuroblasts as indicated by loss of Ase expression and gain of Sp-1::GFP expression. (J) Only type I neuroblasts were found on the wild-type ventral nerve cord. The thoracic segments (Th) are shown. (K) Tll overexpression molecularly transforms type I neuroblasts on the ventral nerve cord into type II neuroblasts. The thoracic segments (Th) are shown. (L) Tll overexpression functionally transforms type I neuroblasts into type II neuroblasts as indicated by the generation of Erm::V5^+^Ase^−^ immature INPs (white arrowheads) and Erm::V5^+^Ase^+^ immature INPs (yellow arrows). Scale bar, 10 μm. Bar graphs are represented as mean ± standard deviation. P-values: *** <0.005. See also Figure S2.

As *tll* is required to maintain type II neuroblasts in an undifferentiated state, direct assessment of *tll*’s role in regulating the competency to generate INPs is not possible. As an alternative, we tested whether Tll overexpression is sufficient to transform type I neuroblasts into type II neuroblasts. To unambiguously distinguish the two types of neuroblasts, we searched for robust protein markers of type II neuroblasts and their progeny. As *Sp-1* mRNA is uniquely detected in type II neuroblasts (Yang et al., 2016), we evaluated Sp1 protein expression. Using an *Sp-1::GFP(BAC)* transgene, we found that Sp1::GFP is expressed in all type II neuroblasts and their progeny, which are exclusively located on the dorsal surface of the brain (Figure 2G). Importantly, Sp1::GFP was undetectable in all type I neuroblasts and their progeny in the brain as well as in the ventral nerve cord (Figures 2G, 2H, 2J). Thus, Sp1::GFP is a new marker for the type II neuroblast lineage (Figure 2A). Many type I neuroblasts ectopically expressing *tll* lost Ase expression but gained Sp1::GFP expression in the brain and the ventral nerve cord (Figures 2I and 2K). Thus, ectopic *tll* expression is sufficient to molecularly transform type I neuroblasts into type II neuroblasts (Figure 2A). Next, we tested if type I neuroblasts ectopically expressing *tll* can generate immature INPs. We examined the expression of Erm::V5, which is expressed in all immature INPs except those that are newly generated (Figure 2A). Indeed, we detected Ase^−^ (Erm::V5^+^Ase^−^) and Ase^+^ (Erm::V5^+^Ase^+^) immature INPs in many type I neuroblast lineages that ectopically express *tll* (Figure 2K; data not presented). These data led us to conclude that ectopic *tll* expression is sufficient to molecularly and functionally transform type I neuroblasts into type II neuroblasts. We conclude that *tll* is a master regulator of type II neuroblast functional identity.

### Ham is a new regulator of INP commitment

*tll* is a putative Notch target in neuroblasts (Zacharioudaki et al., 2016). Because Notch signaling becomes re-activated in INPs, *tll* must become inactivated during INP commitment to prevent supernumerary type II neuroblast formation at the expense of the generation of differentiated cell types (Figure 2A). We predicted that genes required for inactivating *tll* should become upregulated in immature INPs in wild-type and *brat-*null brains following 9 and 18 hours of Insb overexpression. From our RNA sequencing dataset, we identified genes that encode transcription factors and are upregulated in *brat-*null brains overexpressing Insb for 9 or 18 hours. We performed reverse transcriptase PCR and confirmed that the transcript levels of these genes indeed become upregulated in *brat-*null brains overexpressing Insb (Figure 1B; data not presented). Thus, these genes are candidates for inactivating *tll* during INP commitment.

*erm*^*hypo*^ brains are highly sensitive to the changes in gene activity required to prevent INPs from reverting to type II neuroblasts (Janssens et al., 2017). Because ectopic *tll* expression in INPs leads to supernumerary type II neuroblast formation, reducing the function of genes that inactivate *tll* should enhance the supernumerary neuroblast phenotype in *erm*^*hypo*^ brains. We found that *erm*^*hypo*^ brains alone contained 31.5 ± 6.8 type II neuroblasts per lobe (Figure 3A; n = 10 brains). Although reducing the function of most candidate genes did not enhance the supernumerary neuroblast phenotype in *erm*^*hypo*^ brains, knocking down *ham* function with two different *UAS-RNAi* transgenes reproducibly enhanced the phenotype (Figure 3A; n = 10 brains per transgene). Consistent with the effect of reducing *ham* function by RNAi, the heterozygosity of a deficiency that deletes the entire *ham* locus also enhanced the supernumerary neuroblast phenotype in *erm*^*hypo*^ brains (Figure S3A). Thus, *ham* likely plays a role in preventing INPs from reverting to supernumerary type II neuroblasts.

**Figure 3.**
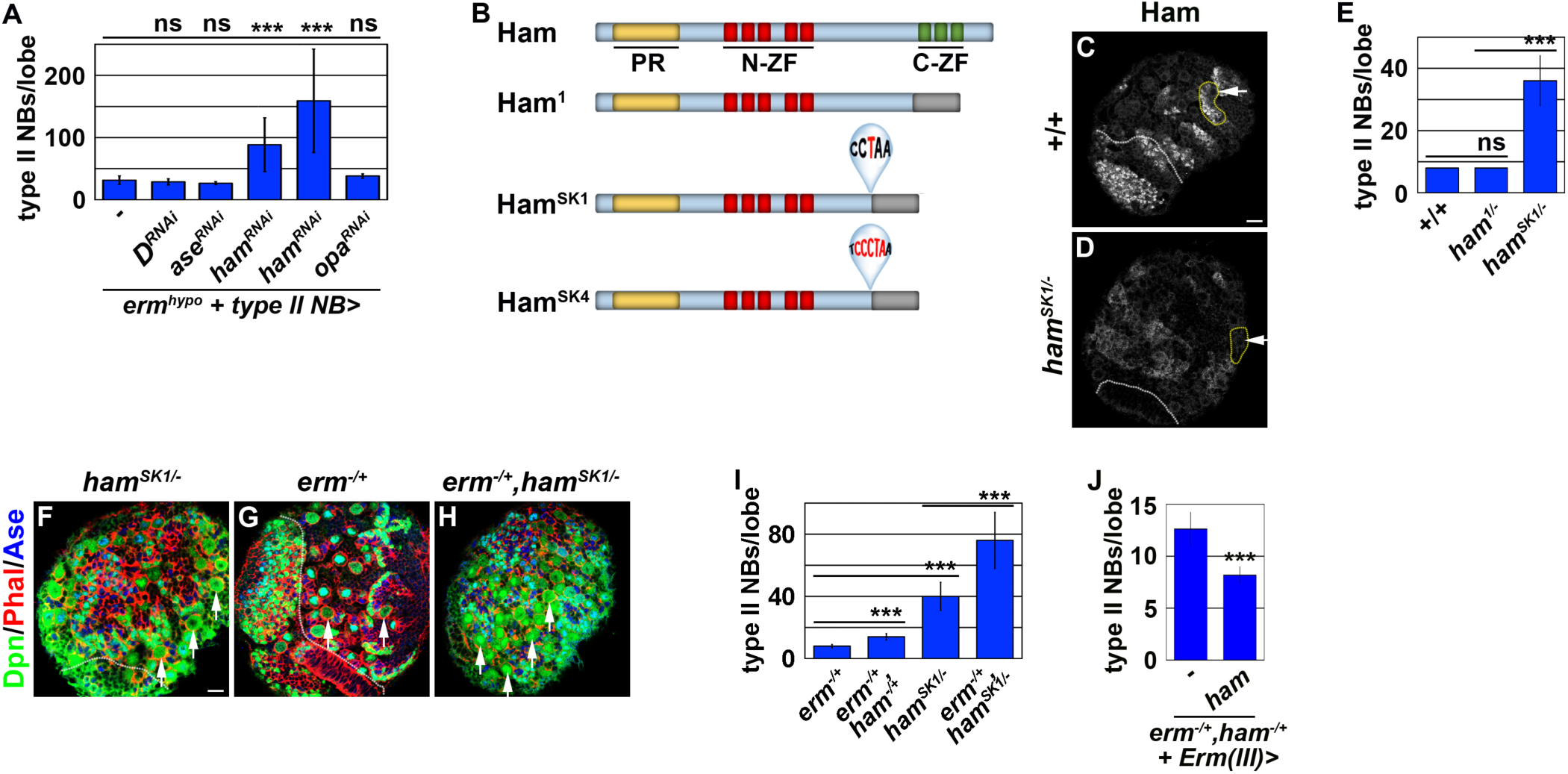
Ham is a novel regulator of INP commitment. (A) Quantification of total type II neuroblasts per lobe in *erm*^*hypo*^ brains with knockdown of candidate regulators of type II neuroblast functional identity. Knocking down *ham* function consistently enhanced the supernumerary type II neuroblast phenotype in *erm*^*hypo*^ brains. (B) A diagram summarizing the lesions in *ham* alleles. (C-D) *ham*^*SK1*^ is a protein-null allele. Ham was detected in immature INPs and INPs in wild-type brains, but was undetectable in *ham*^*SK1*^ homozygous brains. type II neuroblast: white arrows. (E) Quantification of total type II neuroblasts per lobe in *ham* mutant brains. *ham*^*SK1*^ homozygous but not *ham*^*1*^ homozygous brains displayed a supernumerary type II neuroblast phenotype. (F-H) The heterozygosity of *erm* alone had no effect on type II neuroblasts, but enhanced the supernumerary type II neuroblast phenotype in *ham*^*SK1*^ homozygous brains. type II neuroblast: white arrows. (I) The heterozygosity of *erm* enhanced the supernumerary type II neuroblast phenotype in *ham* deficiency heterozygous brains and *ham*^*SK1*^ homozygous brains. (J) Overexpressing Ham in Ase^+^ immature INPs rescued the supernumerary type II neuroblast phenotype in *erm,ham* double heterozygous brains. Scale bar, 10 μm. Bar graphs are represented as mean ± standard deviation. P-values: *** <0.005. ns: not significant. See also Figure S3.

Our findings contradict a previous study that concluded that *ham* does not play a role in suppressing supernumerary type II neuroblast formation (Eroglu et al., 2014). This discrepancy might be due to the *ham*^*1*^ allele used in the previous study, which encodes a nearly full-length protein and exhibits similar protein stability as wild-type Ham (Figure 3B) (Moore et al., 2002). We took two approaches to determine whether *ham* is required for preventing supernumerary type II neuroblast formation. First, we examined *ham* deficiency heterozygous brains and reproducibly observed a mild supernumerary type II neuroblast phenotype (9 ± 0.9 per lobe, n = 12 brains) (Figure S3A). Second, we knocked down *ham* function by overexpressing a *UAS-RNAi* transgene. We found that knocking down *ham* function also led to a mild but statistically significant increase in type II neuroblasts per lobe (9.8 ± 1.8; n = 20 brains) as compared to control brains (8 ± 0; n = 10 brains) (Figure S3B). To confirm that *ham* is indeed required for suppressing supernumerary type II neuroblast formation, we generated two new *ham* alleles, *ham*^*SK1*^ and *ham*^*SK4*^, by CRISPR-Cas9 (Figure 3B). We confirmed that *ham*^*SK1*^ or *ham*^*SK4*^ homozygous brains show undetectable Ham protein expression levels by using an antibody specific for Ham (Figures 3C and 3D; data not presented). We then tested if Ham protein is required for suppressing supernumerary type II neuroblast formation. Indeed, *ham*^*SK1*^ homozygous brains contained 40 ± 10 type II neuroblasts per lobe (Figures 3E and 3F; n = 15 brains). Because Ham is detectable in Ase^+^ immature INPs and remains expressed in all INPs (Eroglu et al., 2014), we conclude that *ham* is a novel regulator of INP commitment.

*ham* knockdown drastically enhanced the supernumerary neuroblast phenotype in *erm*^*hypo*^ brains, leading us to hypothesize that *ham* functions together with *erm* to suppress INPs from reverting to supernumerary type II neuroblasts. We tested this hypothesis by examining a potential genetic interaction between *erm* and *ham* during INP commitment. Although the heterozygosity of *erm* did not increase total type II neuroblasts (8.1 ± 0.3 per lobe; n = 11 brains), it enhanced the supernumerary neuroblast phenotype in *ham* deficiency heterozygous brains (14 ± 2.2 per lobe; n = 14 brains) and in *ham*^*SK1*^ homozygous brains (76.2 ± 18.3 per lobe; n = 14 brains) (Figures 3G-I). We conclude that Ham functions synergistically with Erm to suppress supernumerary type II neuroblast formation.

### Ham promotes stable INP commitment by repressing gene transcription

Re-activation of Notch signaling in INPs drives supernumerary type II neuroblast formation in *erm*-null brains (Weng et al., 2010). Therefore, we hypothesized that supernumerary type II neuroblasts in *ham-*null brains might also originate from INPs. Whereas *erm,ham* heterozygous brains alone contained 12.7 ± 1.6 type II neuroblasts per lobe (n = 9 brains), overexpressing *ham* in Ase^+^ immature INPs and INPs driven by *Erm-Gal4(III)* rescued the supernumerary neuroblast phenotype in *erm,ham* double heterozygous brains (8.2 ± 0.8 neuroblasts per lobe; n = 24 brains) (Figure 3J). This result strongly suggests that supernumerary type II neuroblasts originate from INPs in *ham-*null brains. We generated GFP-marked mosaic clones derived from single type II neuroblasts to confirm the origin of supernumerary type II neuroblasts in *ham-*null brains. In wild-type clones (n = 9 clones), the parental type II neuroblast was always surrounded by Ase^−^ and Ase^+^ immature INPs (Figure 4A). By contrast, supernumerary neuroblasts in *ham*^*SK1*^ homozygous clones (2.5 ± 1.7 type II neuroblasts per clone; n = 11 clones) were always located far away from parental neuroblasts and surrounded by Ase^+^ cells that were most likely Ase^+^ immature INPs and ganglion mother cells (Figures 4B and 4C). It is highly unlikely that supernumerary neuroblasts in *ham*^*SK1*^ homozygous clones originated from symmetric neuroblast division based on their location relative to parental neuroblasts and the cell types that surround them. Thus, we conclude that Ham suppresses INPs from reverting to supernumerary type II neuroblasts.

**Figure 4.**
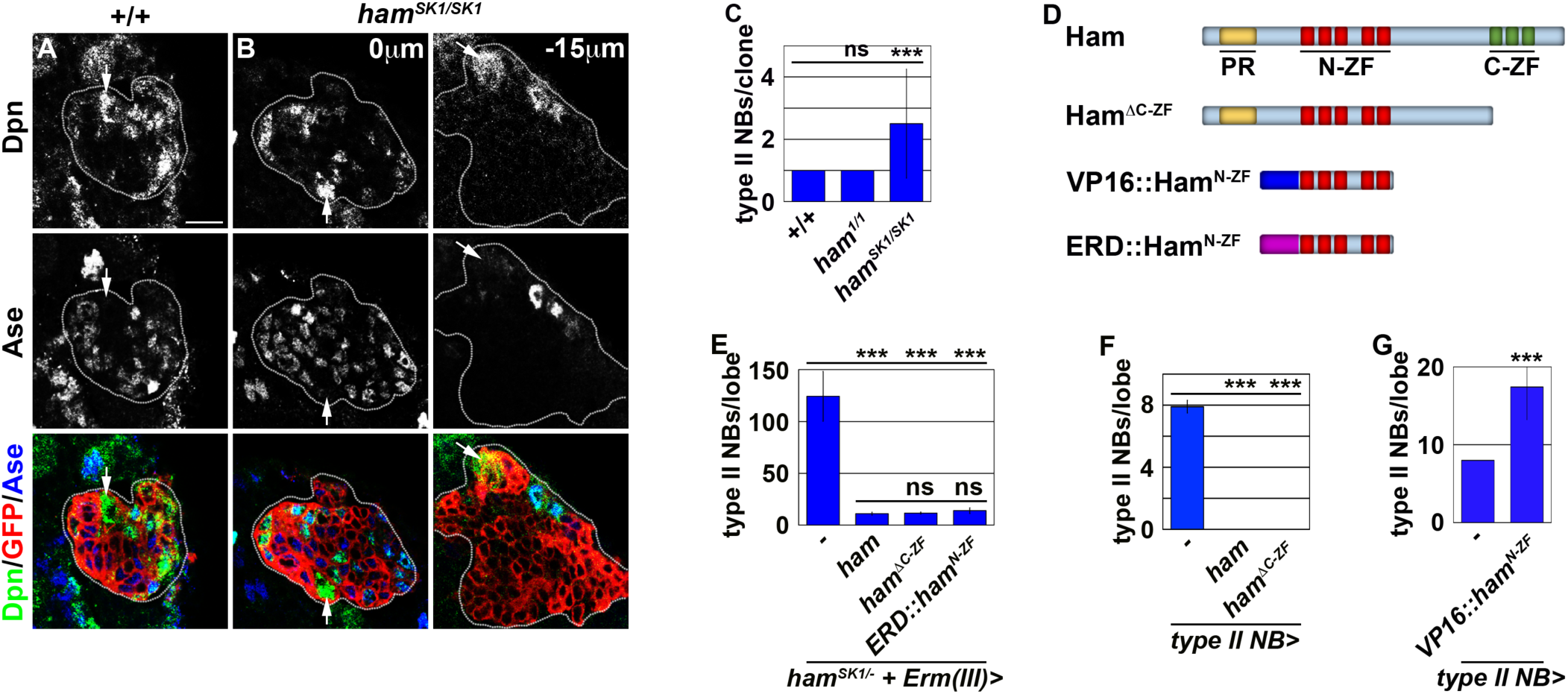
Ham suppresses INP reversion by repressing gene transcription. (A-B) Phenotypic characterization of *ham*^*SK1*^ homozygous type II neuroblast clones. Supernumerary neuroblasts (−15 μm) were always located far from the parental neuroblast (0 μm), and were surrounded by Ase^+^ cells in *ham*^*SK1*^ homozygous clones. type II neuroblast: white arrows. (C) *ham*^*SK1*^ clones contained supernumerary type II neuroblasts but *ham*^*1*^ clones did not. (D) A diagram summarizing *ham* transgenes. (E) Quantification of total type II neuroblasts per lobe in *ham*^*SK1*^ homozygous brains overexpressing various Ham transgenic proteins in Ase^+^ immature INPs. Overexpressing full-length Ham, truncated Ham lacking the C-terminal zinc-finger motif, or ERD::Ham^N-ZF^ rescued the supernumerary neuroblast phenotype in *ham*^*SK1*^ homozygous brains. (F) Brains overexpressing full-length Ham or Ham lacking the C-terminal zinc-finger motif contained fewer type II neuroblasts than control brains. (G) Overexpressing VP16::Ham^N-ZF^ led to supernumerary type II neuroblast formation. Scale bar, 10 μm. Bar graphs are represented as mean ± standard deviation. P-values: *** <0.005. ns: not significant.

Because Erm suppresses supernumerary type II neuroblast formation by repressing gene transcription, Ham likely prevents INP reversion as a transcriptional repressor. We reasoned that Ham might function through its N-terminal zinc finger to prevent INP reversion based on our finding that *ham*^*SK1*^ but not *ham*^*1*^ homozygous clones displayed a supernumerary neuroblast phenotype (Figures 3D and 4C). Consistent with this hypothesis, overexpressing Ham^ΔC-ZF^ rescued the supernumerary neuroblast phenotype in *ham*^*SK1*^ homozygous brains to a similar extent as overexpressing full-length Ham (11.1 ± 1.4 neuroblasts per lobe vs. 11.9 ± 1.3 neuroblasts per lobe; n = 11 or 9 brains, respectively) (Figure 4E). Furthermore, misexpressing full-length Ham or Ham^ΔC-ZF^ triggered premature differentiation in type II neuroblasts (n = 10 brains per genotype) (Figure 4F). These data indicate that Ham functions through the N-terminal zinc-finger motif to prevent INP reversion. Under identical conditions, overexpressing a constitutive transcriptional repressor form of Ham containing only the N-terminal zinc fingers (ERD::Ham^N-ZF^) also rescued the supernumerary neuroblast phenotype in *ham*^*SK1*^ homozygous brains (14 ± 2.3 neuroblasts; n = 9 brains) (Figures 4D and 4E). These results indicate that Ham functions through the N-terminal zinc finger to repress the transcription of genes that can trigger INP reversion to supernumerary type II neuroblasts.

Finally, we tested if the N-terminal zinc finger of Ham mediates target gene recognition. We mis-expressed a constitutive transcriptional activator form of Ham containing only the N-terminal zinc-finger motif (VP16::Ham^N-ZF^) (Figure 4D). We found that VP16::Ham^N-ZF^ misexpression in type II neuroblasts was sufficient to induce supernumerary neuroblast formation (17.1 ± 4.7; n = 10 brains; Figure 4G). Because VP16::Ham^N-ZF^ can exert a dominant-negative effect, we conclude that Ham suppresses INP reversion by recognizing target genes through its N-terminal zinc-finger motif and repressing their transcription.

### Inactivation during INP commitment relinquishes Notch’s ability to activate *tll* in INPs

Our findings strongly suggest that Erm and Ham inactivate *tll* during INP commitment, preventing the re-activation of Notch signaling from triggering *tll* expression in INPs (Figure 5A). Therefore, we tested if Erm and Ham indeed inactivate *tll* by reducing *tll* function in *erm,ham* double heterozygous brains. *tll* heterozygous brains contained 8.2 ± 0.4 type II neuroblasts per lobe (n = 10 brains; Figure 5B), whereas *erm,ham* double heterozygous brains contained 11 ± 1.2 type II neuroblasts per lobe (n = 25 brains; Figure 5B). The heterozygosity of *tll* consistently suppressed the supernumerary neuroblast phenotype in *erm,ham* double heterozygous brains (8.4 ± 0.6 type II neuroblasts per lobe; n = 22 brains; Figure 5B). This result strongly supports our hypothesis that Erm and Ham inactivate *tll* during INP commitment. Consistent with this interpretation, we found that Tll::GFP becomes ectopically expressed in INPs in *erm,ham* double heterozygous brains (Figure 5C). In these brains, we reproducibly observed small cells expressing Tll::GFP and Deadpan (Dpn) but not Ase that were most likely supernumerary type II neuroblasts newly derived from INP reversion (Figure 5D). Furthermore, we found that one copy of the *tll::GFP(BAC)* transgene mildly enhanced the supernumerary neuroblast phenotype in *erm,ham* double heterozygous brains (15.4 ± 1.8 type II neuroblasts per lobe vs. 11.1 ± 1 type II neuroblasts per lobe; n = 12 or 11 brains) (Figure 5E). These data indicate that sequential inactivation by Erm and Ham during INP commitment renders *tll* refractory to activation in INPs.

**Figure 5.**
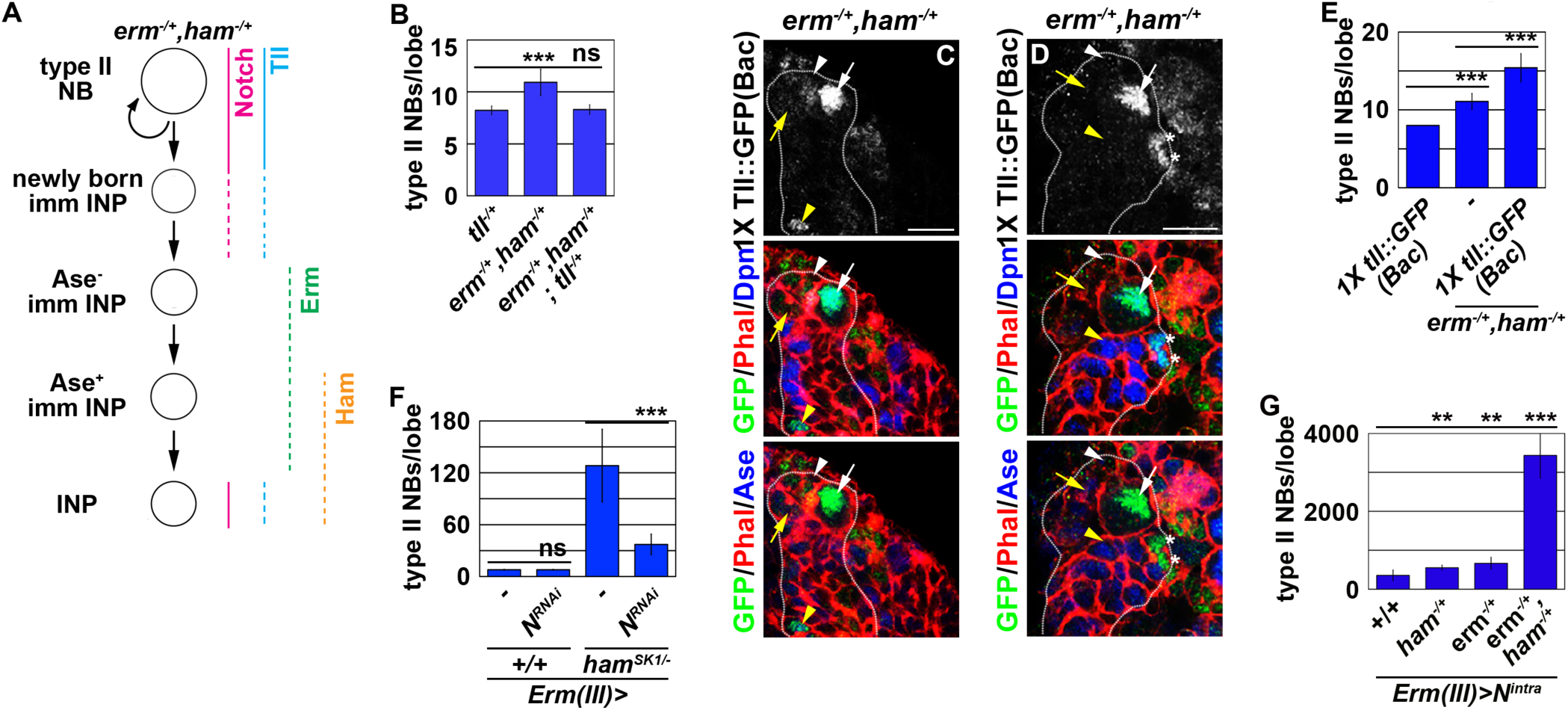
Erm- and Ham-mediated repression renders *tll* refractory for activation by Notch. (A) A diagram depicting our model that ectopic activation of *tll* triggers INP reversion to supernumerary type II neuroblasts in *erm,ham* double heterozygous brains. (B) Heterozygosity of *tll* suppressed the supernumerary neuroblast phenotype in in *erm,ham* double heterozygous brains. (C-D) The *tll::GFP(BAC)* transgene becomes ectopically expressed in INPs and supernumerary type II neuroblasts (*) in *erm,ham* double heterozygous brains. type II neuroblast: white arrows. Ase^−^ immature INP: white arrowhead. Ase^+^ immature INP: yellow arrow. INP: yellow arrowhead. (E) One copy of the *tll::GFP(BAC)* transgene enhanced the supernumerary type II neuroblast phenotype in *erm,ham* double heterozygous brains. (F) Knocking down *Notch* function in INPs suppressed the supernumerary type II neuroblast phenotype in *ham*^*SK1*^ homozygous brains. (G) Overexpressing *N*^*intra*^ induced INP reversion to supernumerary neuroblasts much more efficiently in *erm,ham* double heterozygous brains than *erm* or *ham* heterozygous brains. Scale bar, 10 μm. Bar graphs are represented as mean ± standard deviation. P-values: ** <0.05, *** <0.005. ns: not significant.

Because *tll* is a putative Notch target in neuroblasts (Zacharioudaki et al., 2016), we tested if inactivation of *tll* by Erm and Ham relinquishes the ability of Notch signaling to activate *tll* in INPs. We knocked down *Notch* function in INPs in wild-type or *ham*^*SK1*^ homozygous brains. Knock down of *Notch* function in INPs had no effect on type II neuroblasts in wild-type brains (7.9 ± 0.3; n = 17 brains; Figure 5F). However, in *ham*^*SK1*^ homozygous brains, knocking down *Notch* function in INPs reduced the number of type II neuroblasts per lobe from 128 ± 39.6 (n = 9 brains) to 36.4 ± 11.6 (n = 9 brains) (Figure 5F). This result indicates that re-activation of Notch signaling triggers INP reversion to supernumerary type II neuroblasts in *ham-*null brains. We extended our analyses to test if inactivation by Erm and Ham reduces the competency to respond to Notch signaling. Consistent, overexpressing constitutively activate Notch (Notch^intra^) induced a drastically more severe supernumerary neuroblast phenotype in *erm,ham* double heterozygous brains (3,437.6 ± 586.8 type II neuroblasts per lobe; n = 9 brains) than in *erm* or *ham* single heterozygous brains [687 ± 134.3 (n = 9 brains) and 588.6 ± 77.6 (n = 11 brains) type II neuroblasts per lobe, respectively] (Figure 5G). Together, these data strongly suggest that inactivation by Erm and Ham during INP commitment renders *tll* refractory to Notch signaling in INPs (Figure 5A).

### Inactivation by Hdac3 relinquishes Notch’s ability to activate *tll* in INPs

To prevent Notch signaling from activating *tll* in INPs, Erm and Ham must function through chromatin-modifying proteins. We first tested if Erm and Ham inactivate *tll* by promoting sequential chromatin changes during INP commitment. We overexpressed full-length Ham or Ham^ΔC-ZF^ driven by *Erm-Gal4(III)* in *erm*-null brains. *erm*-null brains alone contained 1,824.6 ± 520.7 type II neuroblasts per lobe (n = 13 brains) (Figures 6A and S4A). Overexpressing full-length Ham or Ham^ΔC-ZF^ strongly suppressed the supernumerary neuroblast phenotype in *erm*-null brains [34.5 ± 8.3 (n = 13 brains) and 64.6 ± 18.3 (n = 7 brains) neuroblasts per lobe, respectively] (Figures 6A and S4B). This result indicates that Ham can replace endogenous Erm function to suppress INP reversion, suggesting that these two transcriptional repressors function through an identical chromatin-modifying protein to inactivate *tll* during INP commitment.

**Figure 6.**
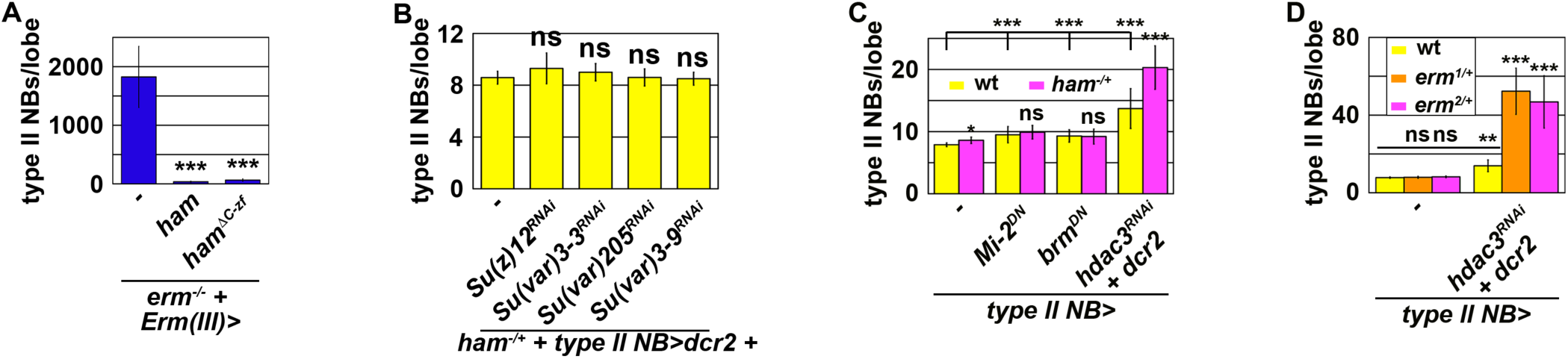
Erm and Ham function through Hdac3 to prevent INPs from reverting to type II neuroblasts. (A) Overexpressing full-length Ham or truncated Ham lacking the C-terminal zinc-finger motif suppressed the supernumerary type II neuroblast phenotype in *erm*-null brains. (B) Reducing activity of PRC2 or the chromatin complex that promotes heterochromatin formation did not enhance the supernumerary neuroblast phenotype in *ham* heterozygous brains. (C) Reducing the activity of Hdac3 but not the Mi-2 or Brm complex enhanced the supernumerary neuroblast phenotype in *ham* heterozygous brains. (D) Reducing the activity of Hdac3 enhanced the supernumerary neuroblast phenotype in *erm* heterozygous brains. Bar graphs are represented as mean ± standard deviation. P-values: ** <0.05, *** <0.005. ns: not significant. See also Figure S4.

We predicted that decreasing the activity of a chromatin-modifying protein required for Ham-mediated gene inactivation during INP commitment should enhance the supernumerary neuroblast phenotype in *ham* heterozygous brains. We knocked down the function of genes known to contribute to the inactivation of gene transcription in *ham* heterozygous brains. Reducing the activity of polycomb-repressive complex 2 (PRC2) or heterochromatin protein 1, as well as decreasing the recruitment of Heterochromatin Protein 1, had no effect on the supernumerary neuroblast phenotype in *ham* heterozygous brains (Figure 6B). While reducing the activity of nucleosome remodelers alone led to a mild supernumerary type II neuroblast phenotype, it did not further enhance the supernumerary neuroblast phenotype in *ham* heterozygous brains (Figure 6C). By contrast, reducing the activity of histone deacetylase 3 (Hdac3) alone resulted in a mild supernumerary type II neuroblast phenotype and significantly enhanced the supernumerary neuroblast phenotype in *ham* heterozygous brains (Figure 6C). These results led us to conclude that Ham likely functions through Hdac3 to prevent INP reversion. We next tested if Erm also functions through Hdac3 to prevent INP reversion. *erm* heterozygous brains alone did not display a supernumerary type II neuroblast phenotype (Figure 6D). Knocking down *hdac3* function in *erm* heterozygous brains led to a more than three-fold increase in supernumerary type II neuroblasts as compared to wild-type brains (Figure 6D). Thus, Erm and Ham likely function through Hdac3 to prevent INP from reverting to supernumerary type II neuroblasts by continually inactivating *tll*.

## Discussion

The expansion of OSVZ neural stem cells, which indirectly produce neurons by initially generating intermediate progenitors, drives the evolution of lissencephalic brains to gyrencephalic brains (Cárdenas and Borrell, 2019; Delaunay et al., 2017; Di Lullo and Kriegstein, 2017). Recent studies have revealed important insights into genes and cell biological changes that lead to the formation of OSVZ neural stem cells (Fujita et al., 2019; Namba et al., 2019). However, the mechanisms controlling the functional identity of OSVZ neural stem cells, including the competency to generate intermediate progenitors, remain unknown. In this study, we have provided compelling evidence demonstrating that Tll is necessary and sufficient for the maintenance of an undifferentiated state and the competency to generate intermediate progenitors in type II neuroblasts (Figure 7). We also showed that two sequentially activated transcriptional repressors, Erm and Ham, inactivate *tll* during INP commitment to ensure normal indirect neurogenesis in larval brains. We identified Hdac3 as the key chromatin-modifying protein that functions with Erm and Ham to prevent aberrant INP reversion to supernumerary type II neuroblasts. We propose that continual inactivation of stem cell functional identity genes by histone deacetylation allows intermediate progenitors to stably commit to generating sufficient and diverse differentiated cells during neurogenesis.

**Figure 7.**
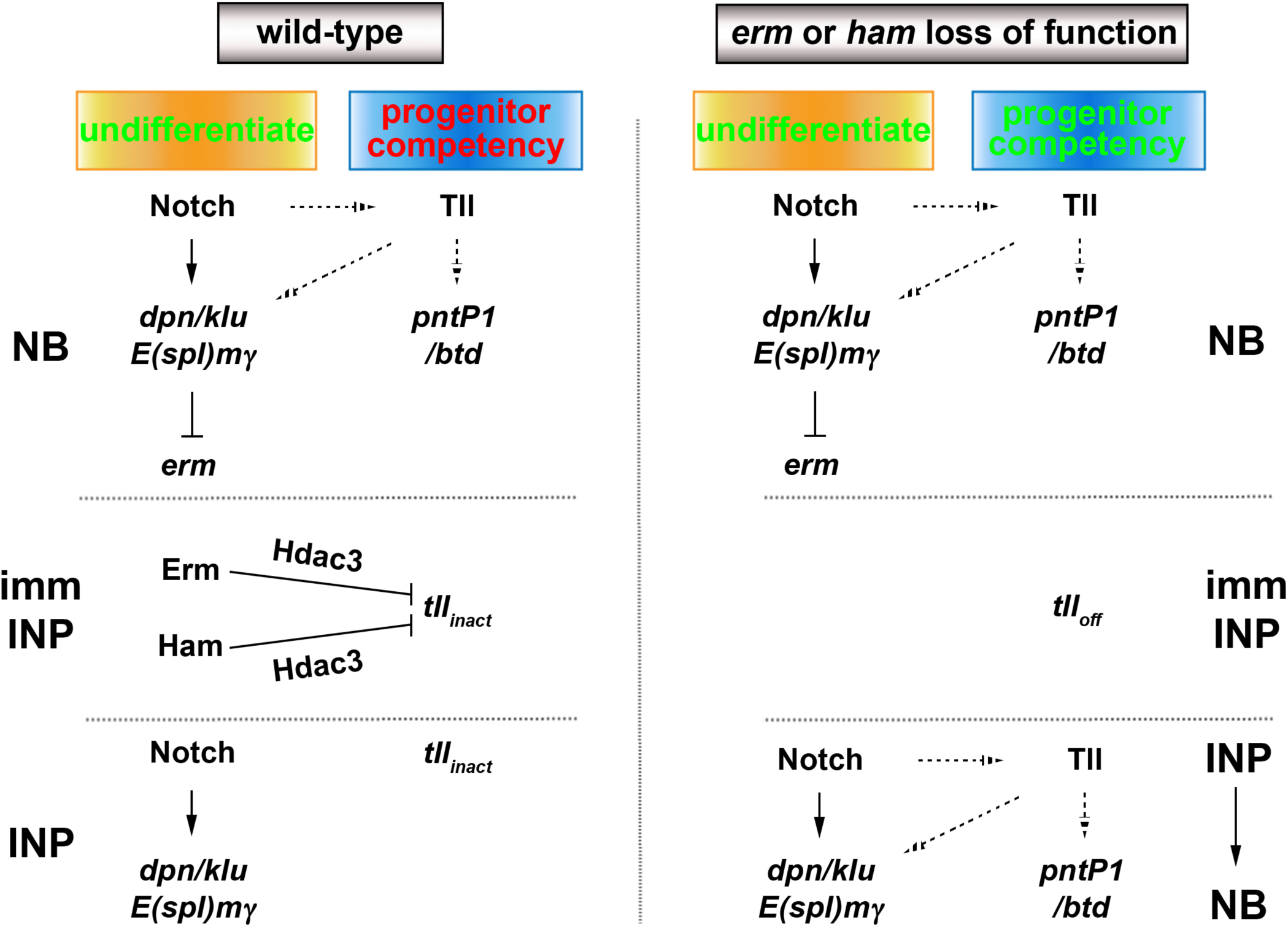
A proposed model for the regulation of type II neuroblast functionality.

### Regulation of the maintenance of an undifferentiated state and the competency to generate intermediate progenitors in neural stem cells

Stem cell functional identities encompass the maintenance of an undifferentiated state and other unique functional features, such as the competency to generate intermediate progenitors. Because genetic manipulation of Notch signaling perturbs the regulation of differentiation during asymmetric stem cell division (Imayoshi et al., 2010; Kageyama et al., 2008), the role of Notch in regulating other stem cell functions remains poorly understood. In the fly type II neuroblast lineage, overexpressing Notch^intra^ or the downstream transcriptional repressors Dpn, E(spl)mγ, and Klumpfuss (Klu) induces the formation of supernumerary type II neuroblasts at the expense of generating immature INPs via the inhibition of *erm* activation (Janssens et al., 2017). Thus, Notch signaling prevents type II neuroblast from initiating differentiation by maintaining *erm* in a poised but inactivated state. By contrast, overexpressing Notch^intra^ but not Dpn, E(spl)mγ, and Klu in combination is sufficient to re-establish a type II neuroblast-like undifferentiated state in INPs, which drives supernumerary type II neuroblast formation. Thus, the Notch-Dpn/E(spl)mγ/Klu axis maintains type II neuroblasts in an undifferentiated state, and Notch alone is a component of the regulatory circuit that endows type II neuroblasts with the competency to generate INPs.

We propose that *tll* is the master regulator of type II neuroblast functional identity (Figure 7). A previous study found that Suppressor of Hairless, the DNA-binding partner of Notch, binds the *tll* locus in larval brain neuroblasts (Zacharioudaki et al., 2016). Identical to *Notch, tll* is necessary and sufficient to induce an undifferentiated state associated with type II neuroblasts. However, overexpressing *tll* but not Notch^intra^ is sufficient to molecularly and functionally transform a type I neuroblast into a type II neuroblast. Thus, Notch likely contributes to *tll* activation, but an additional activator that is specifically expressed in type II neuroblasts must also exist. The transcription factor Zelda (Zld) is expressed in type II neuroblasts but not in INPs and binds the *cis*-regulatory region of *tll* in embryos (Harrison et al., 2011; Nien et al., 2011; Reichardt et al., 2018). However, although all type I neuroblasts exhibit high expression levels of both activated Notch and Zld, Tll is only detectable in a subset of type I neuroblasts (Kurusu et al., 2009; Reichardt et al., 2018). Thus, it is unlikely that Zld plays a key role in activating *tll* expression in type II neuroblasts. The vertebrate homolog of Tll, Tlx, has been shown to function as a transcriptional activator during neurogenesis (Sun et al., 2017). Thus, Tll might function in a positive feedback loop to amplify its own expression in type II neuroblasts. Elucidating the mechanisms that function cooperatively with Notch signaling to activate *tll* expression in type II neuroblasts might provide novel insights into the specification of OSVZ neural stem cells.

### Successive transcriptional repressor activity inactivates stem cell functional identity genes during progenitor commitment

The identification of *ham* as a putative regulator of INP commitment was unexpected given that a previously published study concluded that Ham functions to limit INP proliferation (Eroglu et al., 2014). Ham is the fly homolog of Prdm16 in vertebrates and has been shown to play a key role in regulating cell fate decisions in multiple stem cell lineages (Baizabal et al., 2018; Harms et al., 2015; Moore et al., 2002; Shimada et al., 2017). Prdm16 contains two separately defined zinc-finger motifs, with each likely recognizing unique target genes. Prdm16 can also function through a variety of cofactors to activate or repress target gene expression, independent of its DNA-binding capacity. Thus, Ham can potentially inactivate stemness genes via one of several mechanisms. By using a combination of previously isolated alleles and new protein-null alleles, we demonstrated that the N-terminal zinc-finger motif is required for Ham function in immature INPs. Based on the overexpression of a series of chimeric proteins containing the N-terminal zinc-finger motif, our data indicate that Ham prevents INP reversion to supernumerary type II neuroblasts by recognizing target genes via the N-terminal zinc-finger motif and repressing their transcription. Our results indicate that Ham prevents INPs from reverting to supernumerary type II neuroblasts by repressing target gene transcription.

A key question raised by our study is why two transcriptional repressors that seemingly function in a redundant manner are required to prevent INPs from reverting to supernumerary type II neuroblasts. INP commitment lasts approximately 10-12 hours following the generation of an immature INP; after this time, the immature INP transitions into an INP. *erm* is poised for activation in type II neuroblasts and becomes rapidly activated in the newly generated immature INP less than 90 min after its generation (Janssens et al., 2017). As such, Erm-mediated transcriptional repression allows for the rapid inactivation of type II neuroblast functional identity genes. Because Erm is no longer expressed in INPs when Notch signaling becomes reactivated, a second transcriptional repressor that becomes activated after Erm and whose expression is maintained throughout the life of an INP is required to continually inactivate type II neuroblast functional identity genes. Ham is an excellent candidate because it becomes expressed in immature INPs 4-6 hours after the onset of Erm expression, and is detected in all INPs. Similar to Erm, Ham recognizes target genes and represses their transcription. Furthermore, *ham* functions synergistically with *erm* to prevent INP reversion to supernumerary type II neuroblasts, and overexpressed Ham can partially substitute for endogenous Erm. Thus, Erm- and Ham-mediated transcriptional repression renders type II neuroblast functional identity genes refractory to activation by Notch signaling throughout the lifespan of the INP, ensuring the generation of differentiated cell types rather than supernumerary type II neuroblasts instead.

### Sustained inactivation of stem cell functional identity genes distinguishes intermediate progenitors from stem cells

Genes that specify stem cell functional identity become refractory to activation during differentiation, but the mechanisms that restrict their expression are poorly understood due to a lack of lineage information. Researchers have proposed several epigenetic regulator complexes that may restrict neural stem cell-specific gene expression in neurons (Hirabayashi and Gotoh, 2009; Ronan et al., 2013). We knocked down the function of genes that were implicated in restricting neural stem cell gene expression during differentiation in order to identify chromatin regulators that are required to inactivate type II neuroblast functional identity genes during INP commitment. Surprisingly, we found that only Hdac3 is required for both Erm- and Ham-mediated suppression of INP reversion to type II neuroblasts. Our finding is consistent with a recent study showing that blocking apoptosis in lineage clones derived from PRC2-mutant type II neuroblasts did not lead to supernumerary neuroblast formation (Abdusselamoglu et al., 2019). Our data strongly suggest that genes specifying type II neuroblast functional identity, such as *tll*, are likely inactivated rather than repressed in INPs. This result is supported by the finding that overexpressing Notch^intra^ but not Notch downstream transcriptional repressors in INPs can re-establish a type II neuroblast-like undifferentiated state. We speculate that the chromatin in the *tll* locus remains open and accessible in INPs and that continual histone deacetylation is sufficient to counter the transcriptional activator activity of endogenous Notch and to maintain *tll* in an inactive state (Figure 7). By contrast, the chromatin in the *tll* locus might be close and inaccessible to the Notch transcriptional activator complex; thus, overexpressing Notch^intra^ cannot transform type I neuroblasts into type II neuroblasts. A key remaining question is what transcription factor is required to maintain the chromatin in the *tll* loci in an open state. Thus, insights into regulation of the competency of the *tll* locus to respond to activated Notch signaling might improve our understanding of the molecular determinants of OSVZ neural stem cells.

## Author Contributions

N.Q-R., H.K., and D.H. conducted the experiments. N.Q-R., H.K., D.H., and C.L. designed the experiments. S.K, Q.D., and A.M contributed key reagents. N.Q-R., H.K., and C.L. wrote the manuscript.

## Acknowledgements

We thank Drs. E. Lai and J. Knoblich for providing us with reagents. We thank the Advance Genomics Core and Bioinformatics Core for technical assistance. We thank Dr. Melissa Harrison, Ms. Elizabeth Lawson, and members of the Lee lab for helpful discussions, and Science Editors Network for editing the manuscript. We thank the Bloomington *Drosophila* Stock Center, Kyoto Stock Center, and the Vienna *Drosophila* RNAi Center for fly stocks. We thank BestGene Inc. for generating the transgenic fly lines. This work is supported by NIH grants R01NS107496 and R01NS111647.

## Figure Legends

**Figure S1.**
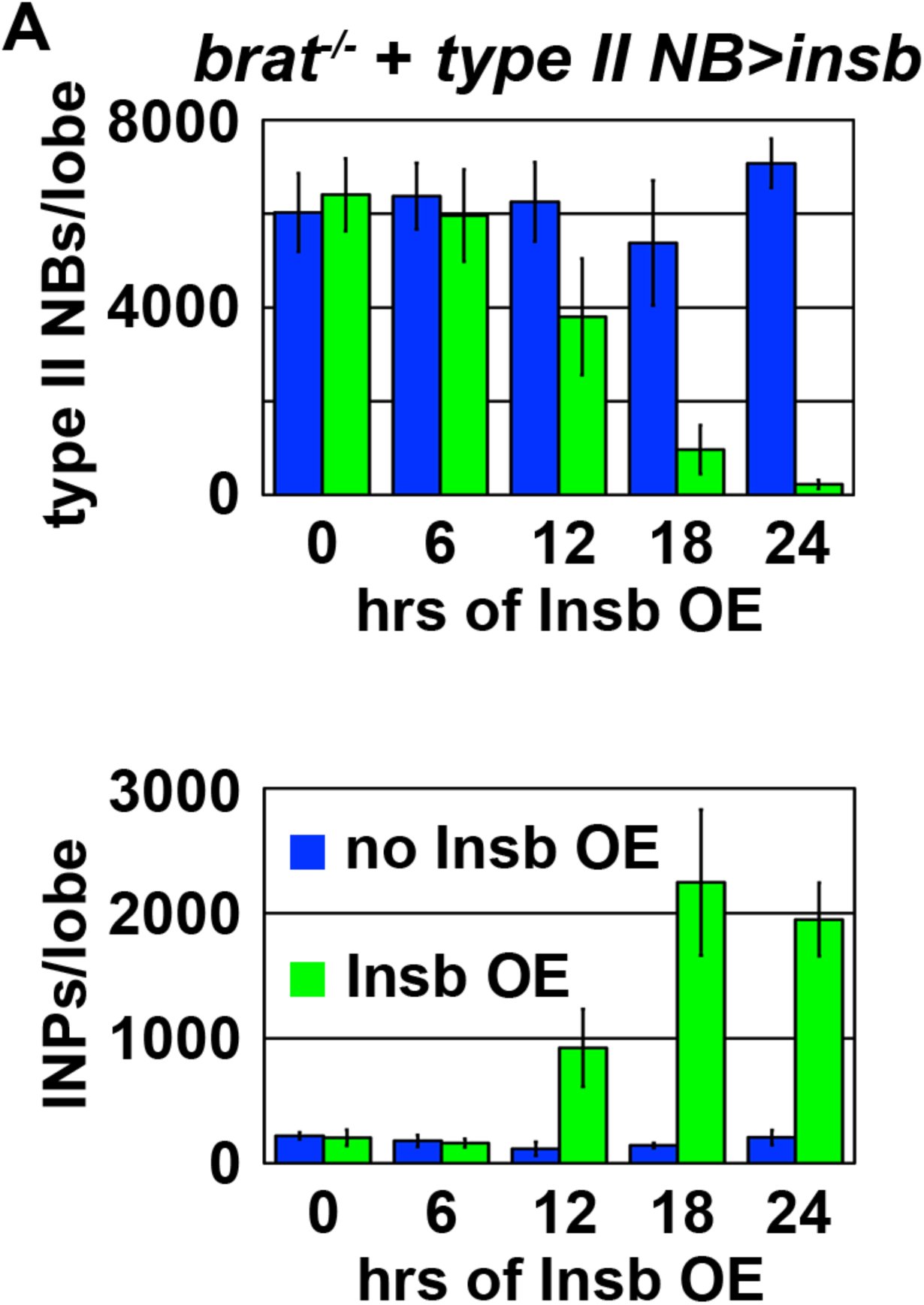
Time course analysis of transient Insb overexpression in *brat*-null brains. Related to Figure 1. Quantification of total type II neuroblasts (top) or INPs (bottom) per lobe in *brat*-null brains following transient Insb overexpression.

**Figure S2.**
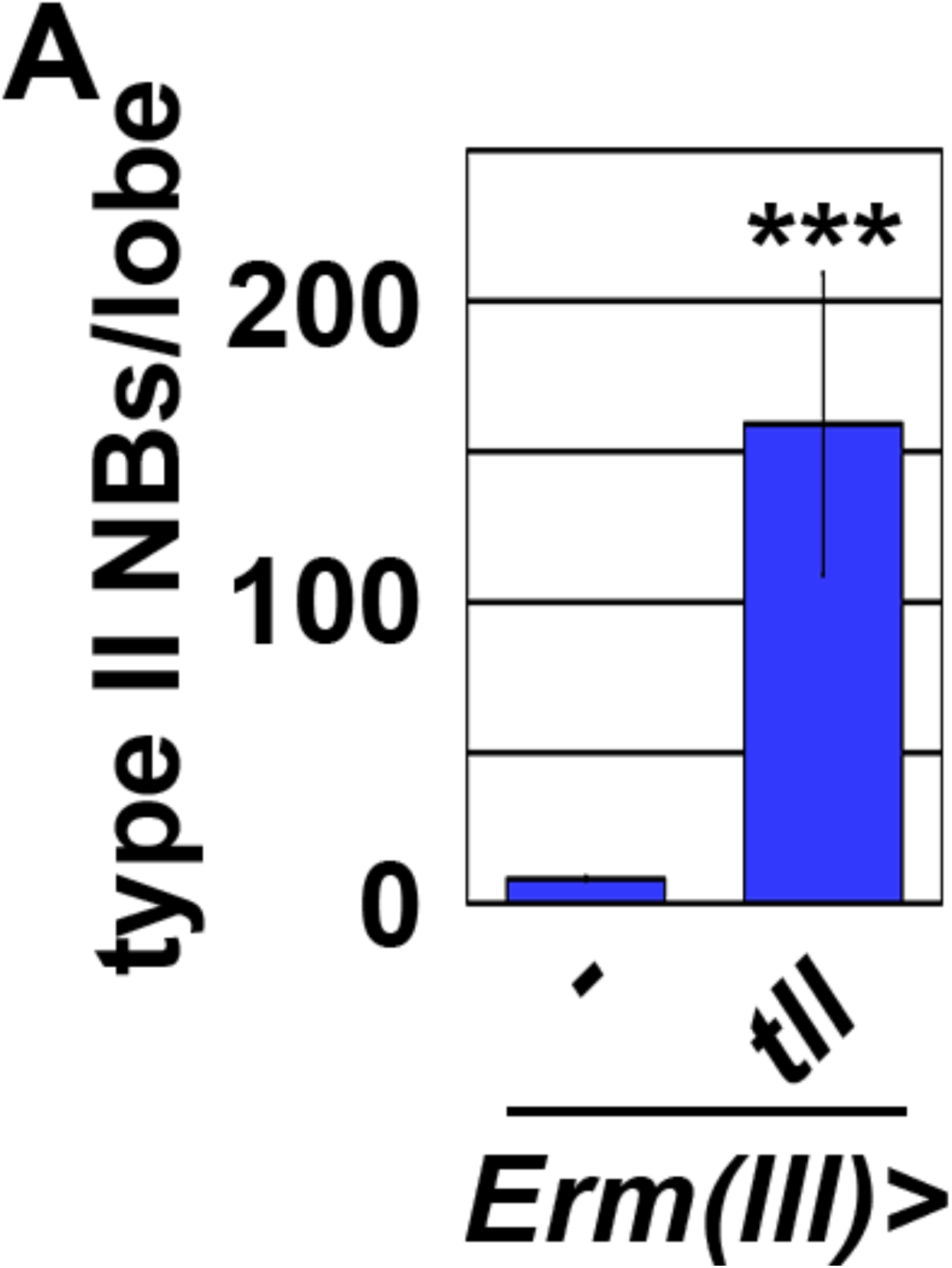
Tll mis-expression in Ase^+^ immature INPs led to supernumerary type II neuroblast formation. Related to Figure 2. Bar graphs are represented as mean ± standard deviation. *** <0.005.

**Figure S3.**
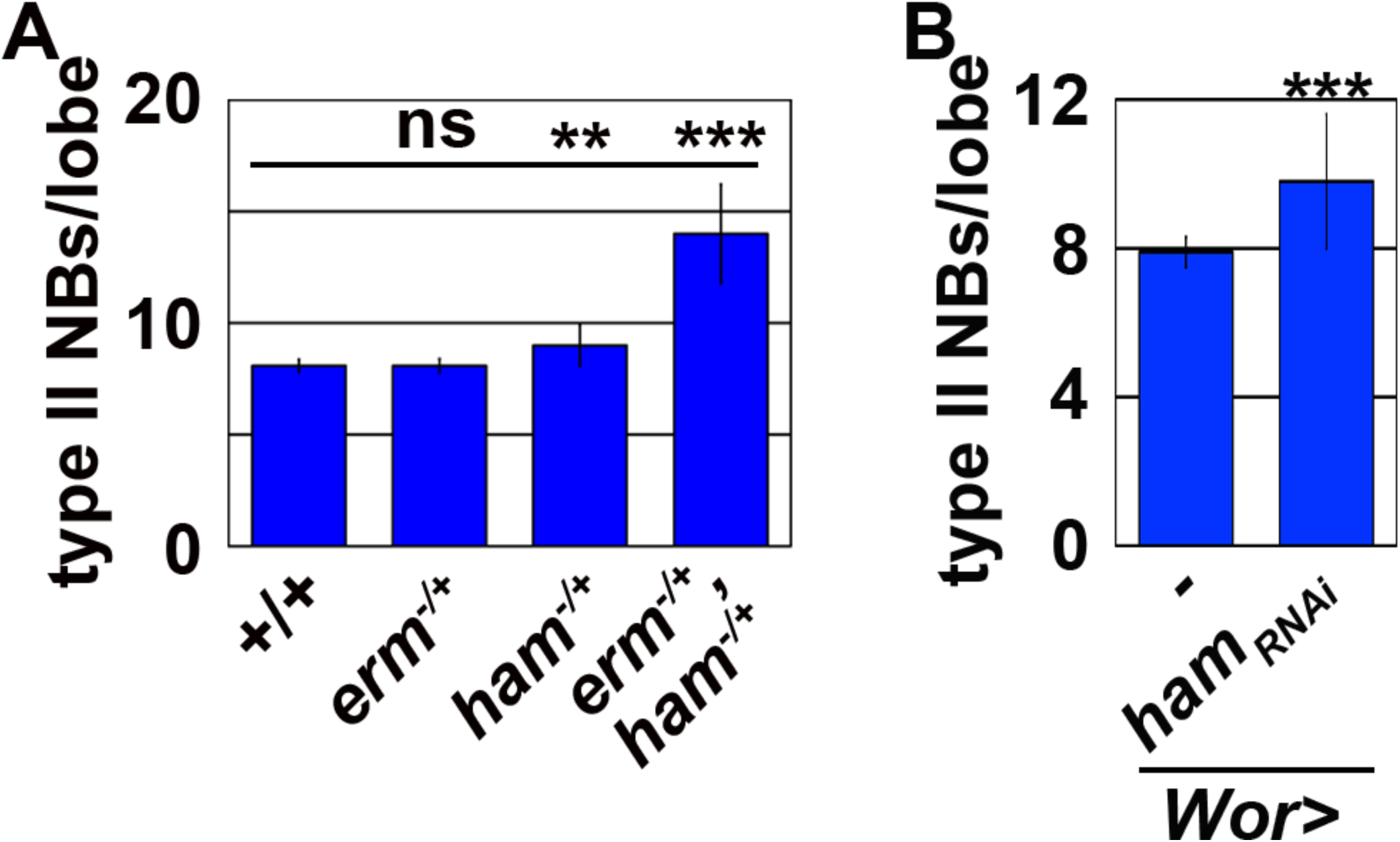
Ham functions synergistically with Erm to suppress supernumerary type II neuroblast formation. Related to Figure 3. (A) *erm,ham* double heterozygous brains displayed the supernumerary type II neuroblast phenotype. (B) Knockdown of *ham* function led to a mild supernumerary type II neuroblast phenotype. Bar graphs are represented as mean ± standard deviation. P-values: ** <0.05, *** <0.005. ns: not significant.

**Figure S4.**
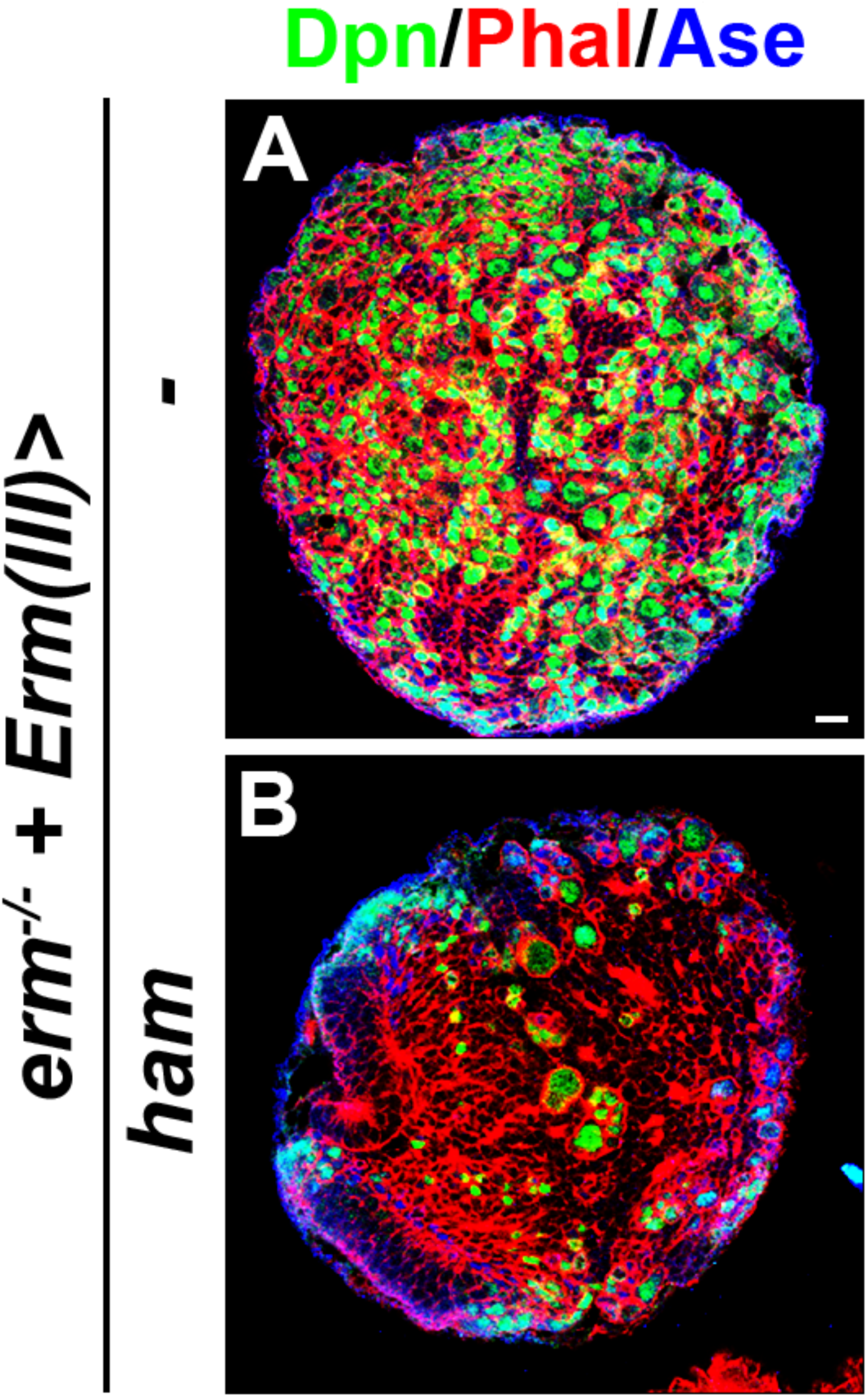
Ham overexpression in Ase^+^ immature INPs suppressed INP reversion in *erm-null* brains. Related to Figure 6. Scale bar, 10 μm.

